# Intron dynamics reveal principles of gene regulation during the maternal-to-zygotic transition

**DOI:** 10.1101/2021.12.01.470832

**Authors:** Kent Riemondy, Jesslyn C. Henriksen, Olivia S. Rissland

## Abstract

The maternal-to-zygotic transition (MZT) is a conserved embryonic process in animals where developmental control shifts from the maternal to zygotic genome. A key step in this transition is zygotic transcription, and deciphering the MZT requires classifying newly transcribed genes. However, due to current technological limitations, this starting point remains a challenge for studying many species. Here we present an alternative approach that characterizes transcriptome changes based solely on RNA-seq data. By combining intron-mapping reads and transcript-level quantification, we characterized transcriptome dynamics during the *Drosophila melanogaster* MZT. Our approach provides an accessible platform to investigate transcriptome dynamics that can be applied to the MZT in non-model organisms. In addition to classifying zygotically transcribed genes, our analysis revealed that over 300 genes express different maternal and zygotic transcript isoforms due to alternative splicing, polyadenylation, and promoter usage. The vast majority of these zygotic isoforms have the potential to be subject to different regulatory control, and over two-thirds encode different proteins. Thus, our analysis reveals an additional layer of regulation during the MZT, where new zygotic transcripts can generate additional proteome diversity.

## INTRODUCTION

During animal oogenesis, the growing oocyte is loaded with maternal RNAs and proteins. After fertilization, the embryo initially is transcriptionally silent, and so the earliest events of development depend on these maternally deposited products. After these initial events, the embryo undergoes the maternal-to-zygotic transition (MZT) and switches developmental control from the maternal genome to the zygotic one (Vastenhouw et al., 2019). The MZT involves two important processes: transcriptional activation of the zygotic genome (known as ZGA for “zygotic genome activation”) and clearance of deposited maternal products, including mRNAs and proteins. Although the timing of these processes differs between species, both are essential for embryonic development (Benoit et al., 2009; Giraldez et al., 2005; Liang et al., 2008).

The fruit fly *Drosophila melanogaster* has played a pivotal role in understanding the molecular choreography of the MZT. In *Drosophila*, as in other species, an initial, minor wave of transcription is followed by more substantial transcriptional activation. Transcription can be detected as early as mitotic cycle 8, and then ~35% of *Drosophila* genes are transcribed in cycles 13–14 (De Renzis et al., 2007; Kwasnieski et al., 2019; Lécuyer et al., 2007; Lott et al., 2011). Possibly due to constraints of rapid cell cycles, initial transcription is associated with short genes, especially those lacking introns, and also with abortive transcription (Edgar and Schubiger, 1986; Heyn et al., 2014; Kwasnieski et al., 2019; Rothe et al., 1992; Shermoen and O’Farrell, 1991). Many of these early transcribed genes contain the so-called “TAG team” motif in their promoter, which is recognized by the pioneer transcription factor Zelda (Liang et al., 2008). Although the continued activity of Zelda through the MZT is required for proper embryogenesis (Liang et al., 2008; McDaniel et al., 2019), its activity is restricted to this developmental window by elegant post-transcriptional gene regulation (Eichhorn et al., 2016; Harrison et al., 2010; Nien et al., 2011).

Extensive research has also defined the *trans*-factors responsible for clearing maternal transcripts during the *Drosophila* MZT. Smaug is a major RNA-binding protein responsible for removal of maternal transcripts. Like Zelda, its mRNA is deposited in the oocyte in a translationally repressed form, and its translation is also upregulated at egg activation (Tadros et al., 2007). Recognizing stem loops in the 3′ untranslated region (3′UTR) of target transcripts, Smaug is responsible for the clearance of nearly one-third of deposited maternal mRNAs (Chen et al., 2014; Tadros et al., 2007). Other trans-factors (e.g., Pumilio (PUM), BRAT, miR-309, and AU-rich element binding proteins) act next to clear many remaining maternal transcripts, including *Smaug* mRNA itself (Bushati et al., 2008; Chen et al., 2014; De Renzis et al., 2007; Laver et al., 2015a; Laver et al., 2015b).

Each of these discoveries has been enabled by approaches classifying genes as maternally or zygotically expressed on a transcriptome-wide scale. There have been several experimental approaches to do so. In zebrafish, RNA polymerase II inhibitors, like a-amanitin, have been injected into early embryos to block ZGA (Lee et al., 2013); although these inhibitors can help determine genes transcribed during initial waves, inhibition of transcription has substantial effects on development and cannot be used beyond this point. In *Drosophila*, strategies have taken advantage of the wealth of genetic and molecular tools. For instance, initial studies made use of chromosomal deletions and microarray analyses to identify zygotically transcribed genes (De Renzis et al., 2007). Another strategy has been to cross two *Drosophila* lines, each with a unique set of single nucleotide polymorphisms (SNPs), to identify zygotically expressed mRNAs, where expression of paternal SNPs is a marker for zygotic transcription (Lott et al., 2011). Finally, a new method is based on injecting the nucleoside analog 5-ethynyl-uridine (5-EU) into embryos prior to ZGA; the analog is incorporated into nascent transcripts, which can then be pulled down by biotinylation of 5-EU through CLICK chemistry and quantified using RNA-seq (Kwasnieski et al., 2019).

However, each method has substantial limitations that can preclude its application to other animal systems. The case of chromosomal deletions is most applicable to *Drosophila melanogaster* and cannot even be readily applied to other *Drosophilids*. Use of SNPs is powerful and can be broadly applied, provided there is a well-characterized genome. However, a major limitation of this approach is the paucity of informative data: the relevant SNP region must be covered with sufficient coverage to accurately quantify allelic expression. For lowly expressed genes, as can be the case for zygotic genes, this requirement can be an issue. Finally, the 5-EU approach requires that embryos be injectable prior to ZGA and that the injection does not alter development. In addition, this approach, while the most sensitive of these methods, becomes less powerful for identifying new transcription of genes after the initial major wave of transcription and may not be well-suited for differentiating between genes with a burst of expression from those with continued expression.

Motivated by these limitations and recent successes in intron-based methods (Gaidatzis et al., 2015; La Manno et al., 2018; Lee et al., 2013; Lugowski et al., 2018), we decided to use intron-mapping reads as a way to identify zygotically transcribed genes. In doing so, we also characterized zygotic transcription at the transcript level, as opposed to just the gene level. To our surprise, these analyses revealed differences in maternal and zygotic isoforms. These differences included different 5′UTRs, coding regions, and 3′UTRs, and were explained by changes in promoter usage, splice isoforms, and cleavage sites, respectively. Maternal and zygotic isoform changes likely affect gene regulation and function. Using published ribosome profiling and mass spectrometry datasets, we found support for expression of both maternal and zygotic protein isoforms, and in some cases, these differences changed protein domains, as is the case for *cycE* and *PNUTS*. Our data suggest that in addition to clearing maternal transcripts and producing zygotic ones, the MZT also changes transcript isoforms and potentially alters gene function. Because our method only requires RNA-seq data from staged embryos and reasonable genome annotation, we propose that it can be applied to characterize the MZT from many systems and be used more broadly to understand other biological processes that involve changes in cell state.

## RESULTS

### Intron signal can be used to define zygotic contributions to the transcriptome

Because introns are spliced out to make mature mRNAs, they are short-lived RNA species and more accurately reflect active transcription than mature transcripts (Gaidatzis et al., 2015; Lugowski et al., 2018). Their instability also makes introns—and intron-mapping RNA-seq reads—low in abundance. But, as library depths have increased, using intron-mapping reads has become possible, enabling a variety of techniques, such as determining cellular “velocities,” normalizing RNA-seq datasets, or identifying gene regulatory programs (Gaidatzis et al., 2015; La Manno et al., 2018; Lugowski et al., 2018). Indeed, analyses of RNA-seq datasets from zebrafish and chicken embryogenesis have previously shown that intron signals aid in defining zygotic transcriptional programs (Hwang et al., 2018; Lee et al., 2013). These approaches employed traditional alignment and quantification strategies that exclude reads that align to multiple locations, which are common for reads derived from repeat rich intronic sequences. We reasoned that quantification approaches that account for ambiguous read alignments would provide an improved method for estimating intronic signal and more accurately define zygotic transcription programs.

We thus developed a pipeline to jointly quantify intron derived and mature RNA species from RNA-seq libraries during embryogenesis (see Materials and Methods for details). In brief, our approach uses the transcript-level quantification tool Salmon (Patro et al., 2017) with a transcriptome reference containing both spliced mRNAs and introns (Figure 1A). Intron sequences are supplemented with adjacent exonic sequences to permit accurate mapping of reads that cross exon-intron junctions. Our pipeline implements a two-pass alignment approach to quantify intron signal. First, libraries from the earliest pre-ZGA samples are aligned to the mRNA-plus-intron transcriptome using Bowtie2. Because the early embryo is transcriptionally silent (Vastenhouw et al., 2019), we reasoned that maternally deposited mRNAs are fully spliced in the fertilized embryo and so intron derived species present reflect ambiguities in transcript annotation. Intronic sequences with detectable read coverage in the pre-ZGA samples are masked to ambiguous nucleotides (N), which prevents productive read-alignment to ambiguous intron sequences. In the second pass alignment, all libraries are realigned to the masked transcriptome and the resulting alignments quantified using Salmon to provide transcript-level abundance estimates for both mature mRNAs and introns. Gene-level or transcript-level estimates of mature mRNA or intronic signals are then aggregated (Soneson et al., 2015). Consistent with our expectations, when we applied our pipeline to published RNA-seq datasets from chicken and zebrafish embryogenesis (Hwang et al., 2018; White et al., 2017), we observed increases in intron signal at developmental stages corresponding to the initiation of zygotic transcription (Figure 1B).

**Figure 1.**
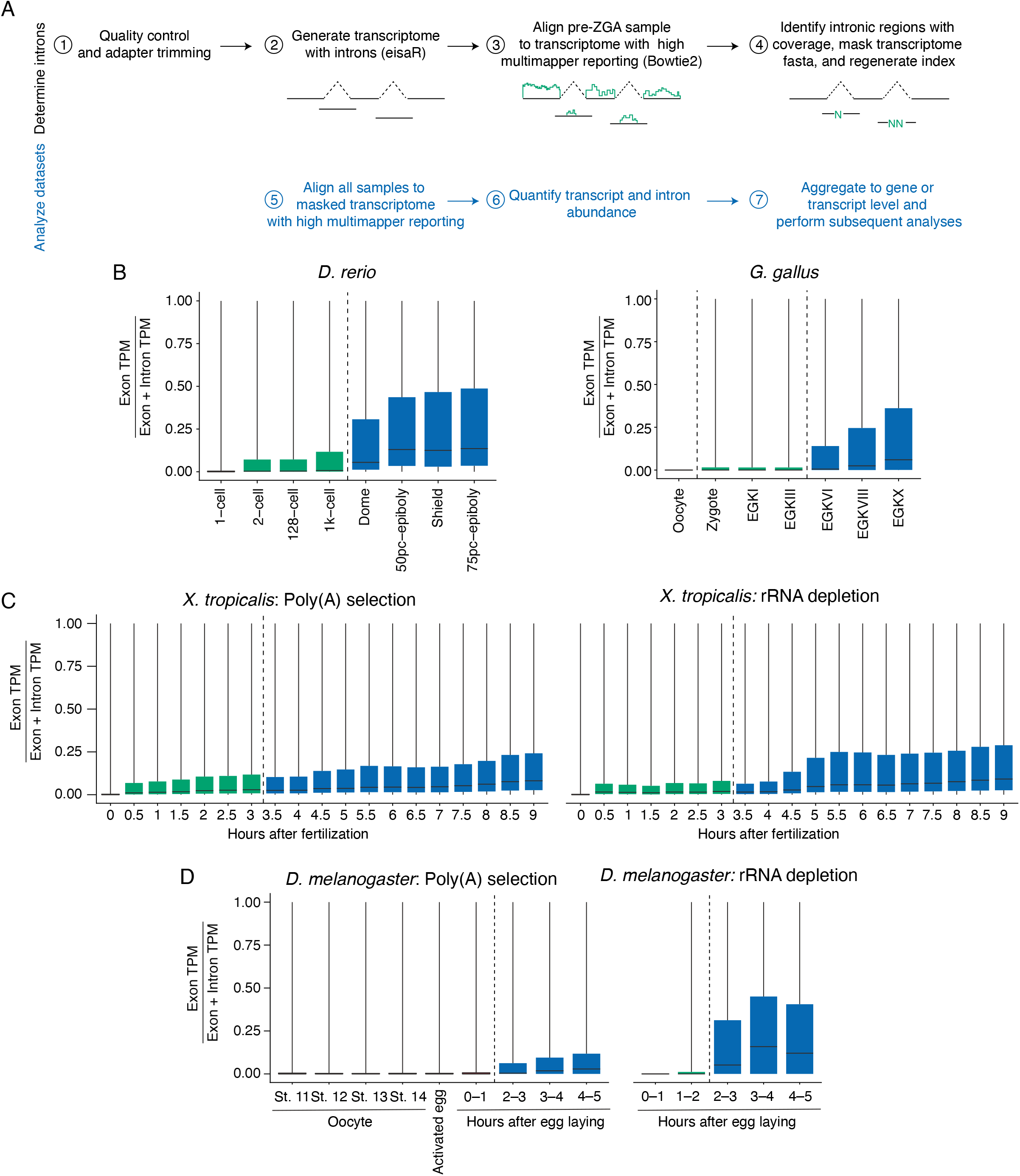
Intron signals are detected at the onset of ZGA. (A) Description of the analytical approach. (B) Quantification of intron signal using the intron masked transcriptome throughout embryonic development in zebrafish (*D. rerio*) and chicken (*G. gallus*). Dashed lines indicate stages of the major waves of de novo zygotic transcription. (C) Quantification of intron signal, as above, throughout *X. tropicalis* embryonic development in either oligo-dT selected or rRNA depleted RNA-seq libraries. (D) Quantification of intron signal, as above, throughout *D. melanogaster* oogenesis embryonic development in either oligo-dT selected or rRNA depleted RNA-seq libraries.

However, one caveat of our initial analyses was that both sets of these libraries were made with oligo(dT) selection. Given that poly(A)-tail length changes during the MZT, we reasoned that extension of poly(A) tails might have confounded our analysis. To investigate this potential issue, we turned to additional datasets from *Xenopus tropicalis* and *Drosophila melanogaster*, comparing RNA-seq libraries produced with oligo(dT) selection and rRNA-depletion (Figure 1C, D). In the case of *Xenopus* (Figure 1C), we analyzed matched oligo(dT)-selected and rRNA-depleted libraries (Owens et al., 2016). In the oligo(dT)-selected dataset, intronic signals increased prior to the known ZGA (~6 hr after fertilization), and no dramatic increase in intron signal was seen at the mid-blastula transition. In contrast, the intron signal increased in the rRNA-depleted dataset at a time point coinciding with ZGA. These results are consistent with cytoplasmic polyadenylation leading to spurious intron quantitation. Moreover, because introns are not polyadenylated, intron-mapping reads have lower abundance in oligo(dT)-selected RNA-seq datasets, meaning that the total RNA dataset has the potential to be more informative for these low abundance RNA species (Figure 1C).

We saw similar results with datasets from *Drosophila* (Figure 1D). As before, we analyzed two datasets. The first dataset contained oligo(dT)-selected libraries from the last few stages of oocyte development (stage 11–14) and the first six hours after egg laying (AEL; 0–1, 1–2, 2–3, 3–4, and 5–6 hr AEL) (Eichhorn et al., 2016). The second dataset contained libraries from the first five hours of embryonic development (0–1, 1–2, 2–3, 3–4, and 4–5 hr AEL), which were made following rRNA-depletion (Wang et al., 2017). Consistent with a lack of transcription during oocyte maturation, we observed no intron signal in the oocyte stages and no or little intron signal in the early embryonic stages prior to the major wave of zygotic transcription for both datasets (Figure 1D). Of note, the earliest stage in each time course was used to generate the intron-masked transcript reference for each dataset (*i.e*., stage 11 from oligo(dT)-selected dataset, and 0-1 hr AEL from rRNA-depleted dataset). In contrast, there was a strong intron signal starting at 2–3 hrs AEL, the time point coinciding with the major wave of zygotic transcription (Figure 1D). For both library preparations, metagene analysis of read coverage surrounding splice sites confirmed that intronic read densities increased at zygotic genes concomitant with the induction of zygotic transcription, but the same was not observed with maternal genes (Supplemental Figure 1). Taken together, we conclude that our pipeline identified hallmarks of zygotic transcription, namely the new production of introns. Oligo(dT)-selected RNA-seq libraries can be used with our pipeline, but using intronic signal to define zygotic expression programs is most appropriate when applied to datasets generated using total RNA species or when applied to organisms that do not have widespread cytoplasmic polyadenylation.

### Dynamics of zygotic transcription in the early *Drosophila* embryo

We next used intron-mapping reads to classify the expression of individual genes as maternal, zygotic, or maternal and zygotic. Because the major wave of transcription occurs 2–3 hours into *Drosophila* development (De Renzis et al., 2007; Kwasnieski et al., 2019; Lécuyer et al., 2007; Lott et al., 2011), we classified libraries from 0–1 and 1–2 hr embryos as containing the maternal RNA complement, while 2–3 hr, 3–4 hr, and 4–5 hr also contained zygotic transcripts. In addition, to avoid confounding effects of changes in poly(A) tail lengths, we classified genes using rRNA-depleted RNA-seq datasets (Wang et al., 2017). Genes with >1 TPM in maternal stages were classified as maternal and those with >2 fold increase (*p_adj_* < 0.05) in either exonic or intronic signal in zygotic stages were considered to have evidence of zygotic transcription, resulting in 1733 maternal genes, 2329 maternal/zygotic, and 1020 zygotic-only genes (Figure 2A; Table S1). Exonic signal alone was able to classify 1637 (48.9%) of genes as zygotically transcribed, whereas intronic signal enabled identification of an additional 1712 genes.

**Figure 2.**
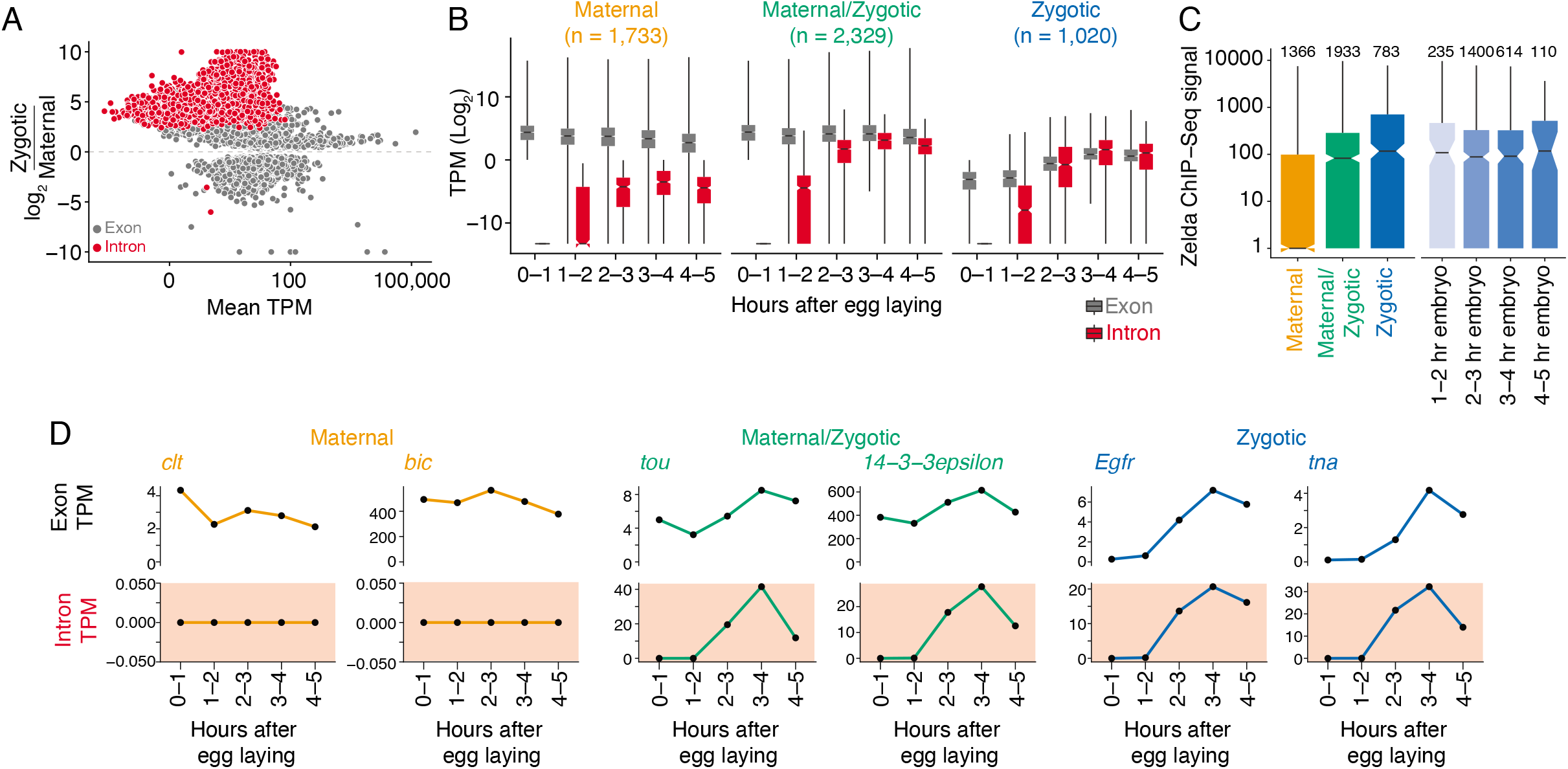
Intron signal identifies the zygotic transcriptome. (A) Comparison of intronic and exonic fold-changes between maternal stages (0-1 and 1-2 hr AEL) and zygotic stages (2-3, 3-4, 4-5 hr AEL). (B) Distribution of TPMs for genes classified as zygotic (<1 TPM in maternal stages, 2-fold increase in exonic or intronic signal with adjusted p-value < 0.05), strictly maternal (>1 TPM in maternal stages, <1 TPM maximal intron signal, adjusted p-values > 0.05 for exonic and intronic signals), or maternal/zygotic (>1 TPM in maternal stages, 2-fold increase in exonic or intronic signal with adjusted p-value < 0.05). (C) Zelda ChIP-Seq signal for each classification (left panel) or for maternal/zygotic and zygotic genes binned by the time to first detectable expression (see methods). (D) TPM distribution for example genes from each category.

Consistent with our classifications, transcripts from zygotic genes were almost 100-fold less abundant than those from maternal genes in the initial time point (Figure 2B). Similarly, maternal genes had low intron signals throughout the time course, while both maternal/zygotic genes and zygotic genes showed an increase in signal at the time point coinciding with ZGA. We note that many maternal genes had detectable increases in intronic signal, but these did not achieve significance. As further evidence of correct classifications, Zelda ChIP-signal (Harrison et al., 2011) was substantially higher for zygotic genes than maternal ones (median 117.05 vs 1; adjusted *p*-value < 5 x 10^-44^), while maternal/zygotic genes showed an intermediate level of Zelda binding (Figure 2C).

We next looked at the relationship between intronic and exonic signals through the MZT. In general, as expected we observed that intronic signal preceded increases in exonic signal (Figure 2D, Supplemental Figure 2). Interestingly, our approach can be used to identify genes undergoing zygotic transcription even when the transcripts are undergoing degradation, as seems to be the case with *14-3-3epsilon*. This result highlights the power of an intron-based method in comparison with other methods (such as SNP calling) that can be affected by mRNA decay.

### A comparison of different methods to annotate the zygotic transcriptome

We next compared our classification of zygotically transcribed genes with previously published methods. First, we looked at our approach in comparison to a traditional intron-counting method (which employs alignment to the genome, followed by counting of uniquely aligned reads overlapping intron features (see Materials and Methods for details). Our approach identifies most zygotic genes detected by the traditional method (73%, 1,646 genes), while detecting an additional 1,703 genes (Figure 3A). Zygotic genes that were detected only by the unique alignment strategy were also significantly differentially expressed in our pipeline, however these genes largely had TPM estimates below our abundance cutoff (>1 TPM) and therefore were not classified (Supplemental Figure 3).

**Figure 3.**
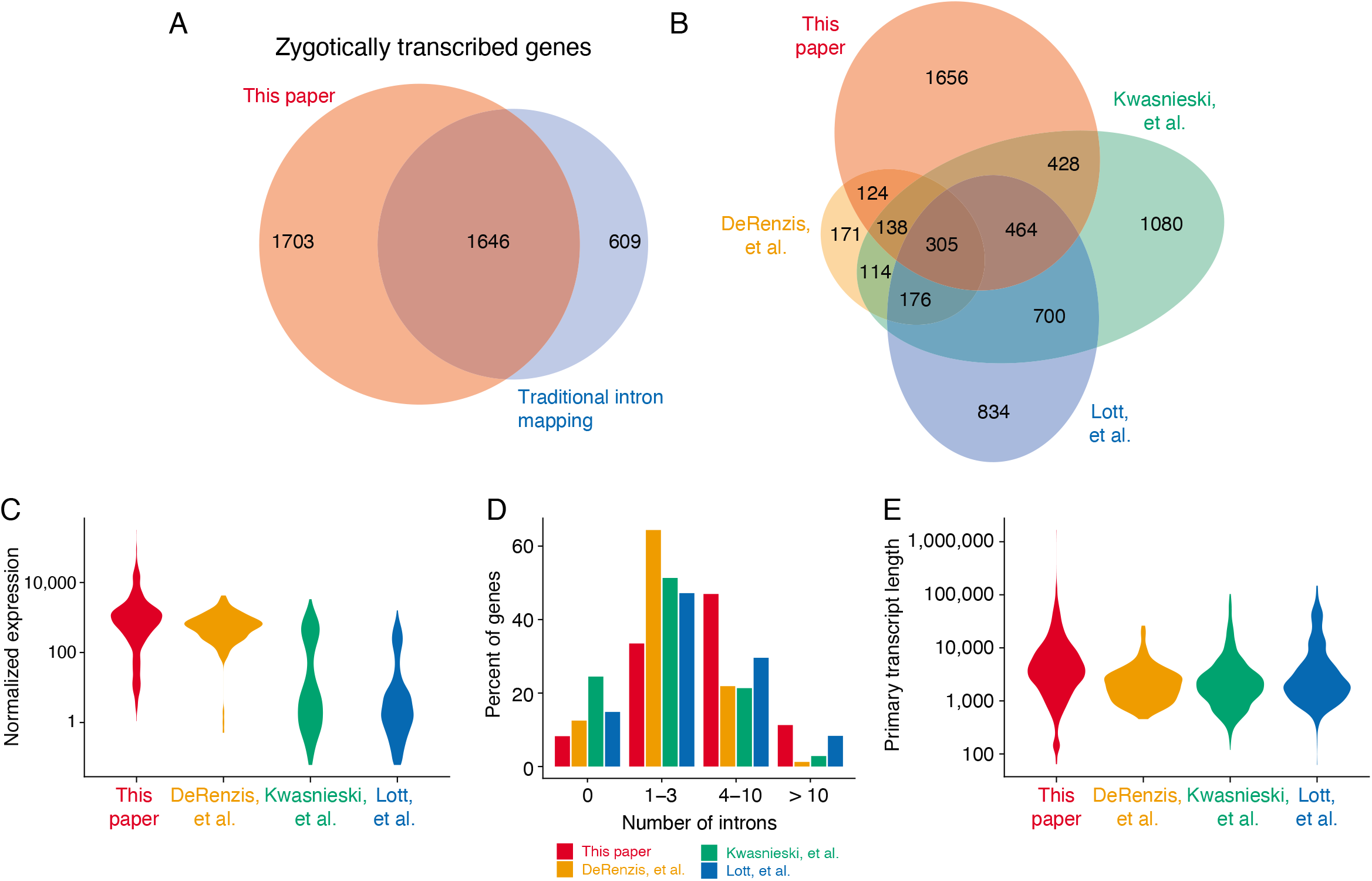
Intron mapping method is more sensitive than traditional alignment and quantification strategies. (A) Comparison of zygotic genes identified by our pipeline and those from application of a traditional alignment and quantification strategy (see methods). (B) Comparison of zygotic genes identified to published experimental approaches. (C) DESeq2 normalized expression of zygotic genes uniquely identified in each study. (D) Number of introns in genes uniquely identified in each study and (E) mean primary transcript length of genes unique to each study.

Next, we examined the specificity of our method by comparing zygotic genes identified via intronic signals to gold-standard classifications generated by orthogonal experimental approaches. Three datasets were examined: a zygotic gene detection via hybrid SNP mapping (Lott et al., 2011), a chromosome deletion approach (De Renzis et al., 2007), or a 5-EU pulldown approach (Kwasnieski et al., 2019). Interestingly, only 481 zygotic genes were detected across all published studies, and between these three experimental methods, there were substantial differences in the genes detected (Figure 3B). Of those genes detected in all three studies, 63% (305) were detected with our approach as opposed to 44% (211) from traditional alignment approaches (Supplemental Figure 4). These results demonstrate that our pipeline provides additional sensitivity to detect zygotic transcription compared to traditional alignment and quantification approaches.

To understand the differences in classification between the different methods and our pipeline, we examined possible explanations. Because introns generally have low abundance, poor overall gene expression can mean that introns could have almost undetectable expression. Consistent with this explanation, genes identified solely by the 5-EU or SNP method had lower expression (Figure 3C). Another obvious requirement for our pipeline is the presence of introns. Indeed, in genes that were classified as zygotically transcribed using our method as opposed to other published methods, the number of introns was larger, and thus overall gene length longer (Figure 3D, E). Taken together, our analyses demonstrate the power of our pipeline over straight intron-counting methods, but there are caveats with it as with any method (see Discussion).

### Identification of dynamic transcript isoforms during the *Drosophila* MZT

Genes whose transcripts are both maternally deposited and zygotically transcribed are the largest class of genes. During our analysis, we noted that several genes, such as the adenosine deaminase *Adar* and the cell cycle regulator *Cdk2*, expressed obviously different maternal and zygotic transcript isoforms based on visual inspection of genome browser tracks (Figure 4A). In the case of *Cdk2*, the zygotic transcript contained an extended 3′UTR. For *Adar* and *Zn72D*, additional coding exons were included in the zygotic isoform, while in the case of *bl* (also known as *hnrnp-k)*, the use of a different promoter led to a different 5′UTR (Figure 4A).

**Figure 4.**
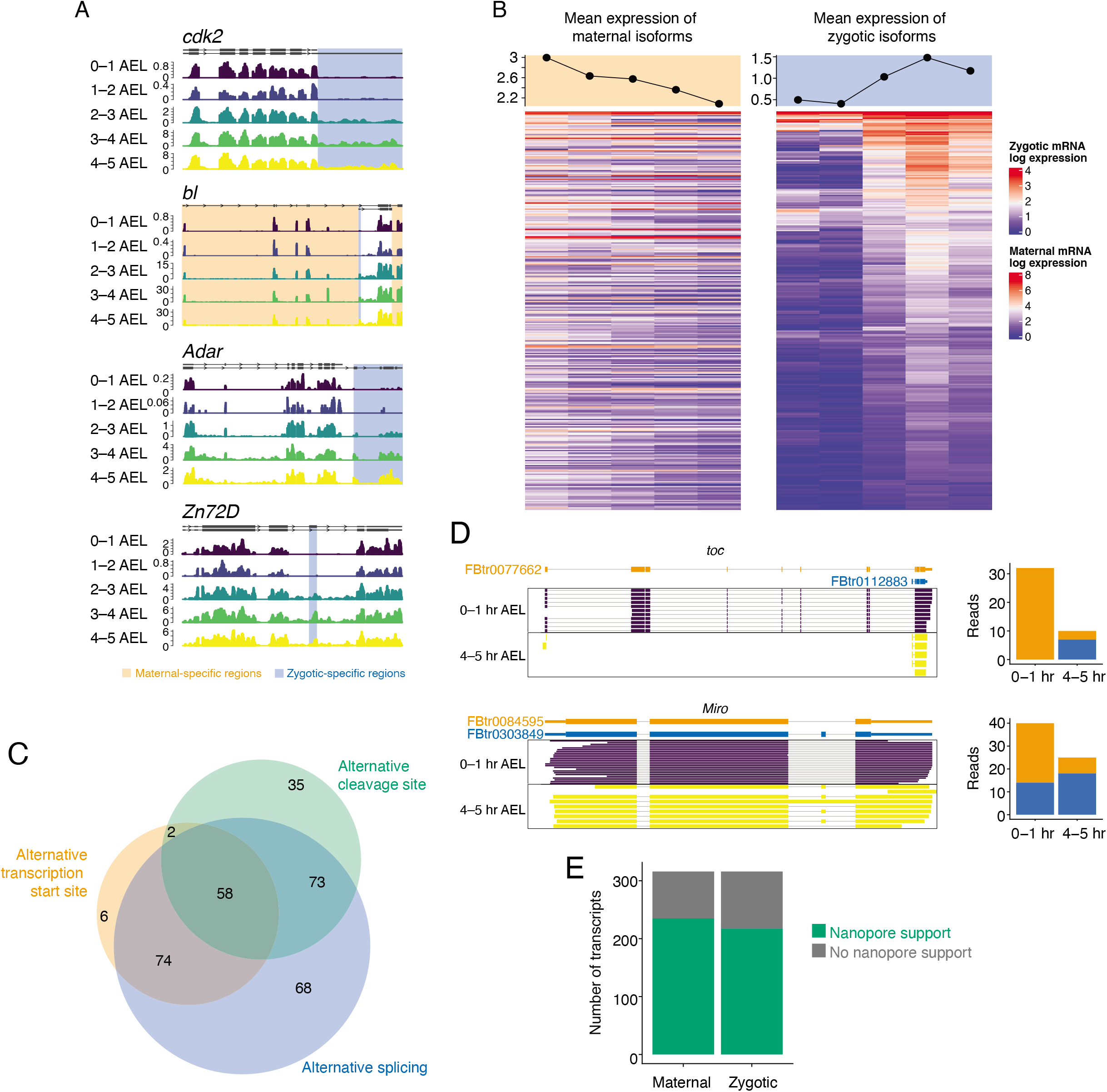
Identification of transcript isoform switching events during ZGA. (A) Read coverage plots for selected genes identified to have an isoform switching event. (B) Expression of the maternal and zygotic isoforms (316 genes) throughout the embryonic time course. (C) Euler diagram depicting alterations in transcript features between maternal and zygotic exons. (D) Read coverage of maternal (orange) and zygotic (blue) transcripts from Nanopore direct RNA-sequencing data. Only mostly full-length (>75%) reads are shown for clarity with quantification of all aligned reads shown in companion bar plots. (E) Quantification of the number of isoforms supported by long-reads in the respective maternal (0-1 hr) or zygotic (4-5 hr) samples.

To explore how general this observation was, we determined changes in isoform usage, comparing transcript-level quantification (derived from exonic sequences) between maternal-only stages (0-1, 1-2 hr AEL) and zygotic stages (2–3, 3-4, 4–5 hr AEL). We used the DEXseq package to identify changes in differential transcript usage, requiring a gene-level p-value less than 0.05, then further filtering to identify genes with both a maternal and zygotic transcript with significant alterations in transcript usage (adjusted p-value < 0.05), and requiring the zygotic isoform to account for at least 10% of the expressed transcripts from a gene.

Using these criteria, we identified 483 genes with alterations in transcript isoforms during the ZGA (Table S2). (Note that in some cases the switch is not binary—*e.g*., a low level of the “zygotic” isoform can be maternally deposited.) Importantly, because the original RNA-seq datasets were made using rRNA-depletion, these results cannot be explained by changes in poly(A)-tail length during the MZT. As expected, many of these genes (316 or 65.4%) were classified by our pipeline as maternal and zygotic, with 153 maternal genes (31.7%) and 14 zygotic-only genes (2.9%). The increase in abundance of zygotic isoforms coincided with the major wave zygotic transcription, while the mean abundance of maternal transcripts decreased during the MZT, consistent with these isoforms being cleared (Figure 4B).

Zygotic transcript isoforms were generated through multiple mechanisms, including alternative splicing, alternative promoter usage, and alternative polyadenylation (Figure 4C). Interestingly, most genes showed a combination of changes. For instance, of the 138 genes that had a different zygotic promoter, 132 also showed differences in alternative splicing and, of those, 58 also had a different transcription termination site. To determine whether individual zygotic transcripts displayed these complex changes, we applied Oxford nanopore technology (ONT) to RNA isolated from 0–1 and 4–5 hr embryos (Figure 4D, E). In the case of *toc*, which is involved in mitotic spindle organization (Debec et al., 2001), the zygotic isoform uses a downstream promoter, resulting in a distinct 5′ UTR and shortened coding region, and we found that this transcript was only produced after ZGA, coinciding with a decrease in the longer maternal isoform. In the case of *Miro*, which is involved in mitochondrial transport on microtubules (Guo et al., 2005), the zygotic isoform includes an additional coding exon and is expressed to some extent prior to ZGA but its levels increase after zygotic transcription; the maternal isoform levels drop during the MZT, presumably due to mRNA decay. In total, although long-read sequencing lacks the depth associated with short-read, Illumina technology, we were able to detect the majority of maternal and zygotic isoforms (75% and 62%, respectively; Figure 4E).

Genes with switching isoforms are enriched in diverse biological processes and molecular activities (Table S3). Notably, many RNA binding proteins displayed switching isoforms, including genes involved in regulation of RNA splicing (including *bl, Hrb27C,pUf68, lark, Rbp1, sqd*), genes involved in dosage compensation and sex determination (*e.g*., *tra2, mle, Zn72D*), double-stranded RNA binding proteins (*e.g., loqs, Adar, DIP1*). and regulators of translation (*e.g*., *eIF-2gamma, tyf, larp, Fmr1, 4EHP, brat*). Other pathways enriched include genes in the cyclin-dependent protein kinase holoenzyme complex (*Cdk2*, *CycE*, *CycT*, *CycA*). Together, these results suggest that, even for genes maternally expressed, zygotic transcription may be an important force shaping gene function.

### Isoform switching generates diversity in the zygotic proteome

We next asked how zygotic isoforms affected coding and regulatory potential of the transcripts. Consistent with previous experiments in zebrafish and *Xenopus* (Owens et al., 2016; Ulitsky et al., 2012), 56% of zygotic isoforms had different 3′UTRs, and 96% had altered 5′ and/or 3′UTRs (Figure 5A). For instance, in the case of *toc* (Figure 4D), use of the downstream promoter likely affects translation: the 5′UTR of the maternal isoform is nearly 440 nucleotides and has three upstream AUGs and a weak Kozak sequence, while that of the zygotic transcript is only 64 nucleotides and has no upstream AUGs and a stronger Kozak sequence. These results highlight the potential for maternal and zygotic transcripts to be subject to different regulation in translation or mRNA decay.

**Figure 5.**
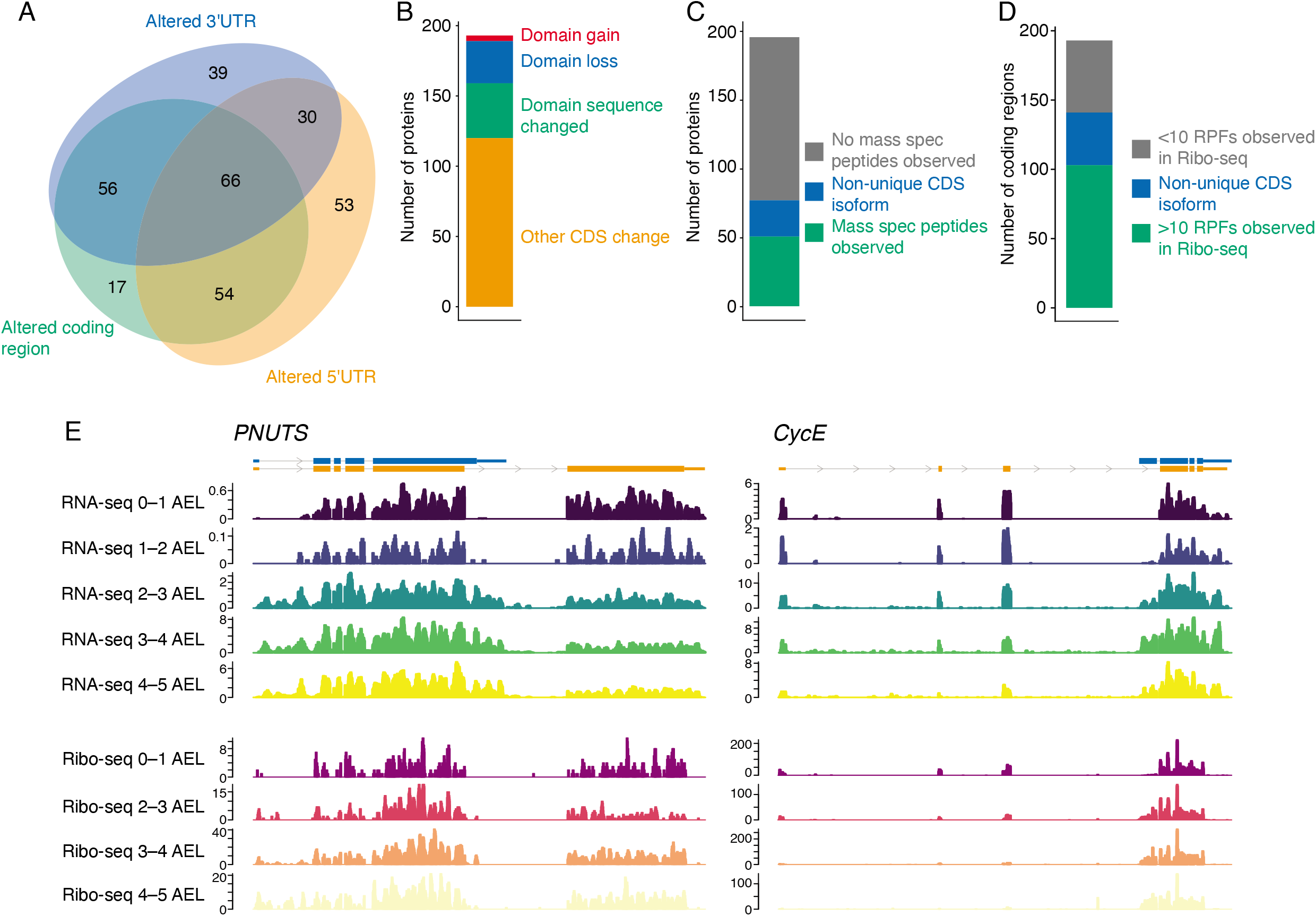
Isoform switching events alter transcript features and coding sequence. (A) Euler diagram depicting alterations in untranslated or CDS regions. Non-coding isoforms are excluded from this analysis. (B) Quantification of the impact of zygotic CDS sequence changes on protein domains (PFAM) or CDS sequence. (C) Quantification of zygotic CDS sequences supported by the presence of peptides in mass spec data from embryonic samples (0 – 6 hours AEL). CDS regions that are not distinguishable between maternal and zygotic are indicated as non-unique CDS isoforms (e.g., due to a truncated CDS). (D) Quantification of zygotic CDS regions with at least 10 ribosome protected fragments (RPFs) detected from ribosome profiling experiments. (E) RNA-seq read and Ribosome profiling RPF coverage across selected genes that undergo isoform switching resulting in alterations in CDS sequence.

We were intrigued to note that 67% of the zygotic isoforms are also predicted to lead to altered proteins, as we saw with *Miro* (Figure 4D). Many of these changes likely alter protein function. For instance, as mentioned above, *Adar*, which encodes the adenosine deaminase responsible for A-to-I editing, has different maternal and zygotic isoforms. The maternal transcript is shortened (FBtr0305497) and encodes a truncated deaminase domain lacking 272 amino acids, which is restored in the dominant zygotic isoform (FBtr0307895). Our observation is consistent with previous RT-PCR results (Ghosh et al., 2013). Interestingly, A-to-I RNA editing is known to increase during embryonic development (Graveley et al., 2011), and it is tempting to speculate that this result is in part due to zygotic production of the longer isoform. Similarly, the zygotic isoform of *bl* lacks two of the three KH RNA binding domains. Overall, 30 (15.5%) of the coding regions in the zygotic isoforms had lost domains, while four (2.0%) gained domains or 39 (20.2%) had sequence alterations to domains (Figure 5B).

We next asked if there was evidence of these zygotic isoforms producing proteins. First, we examined publicly available mass spectrometry data (Casas-Vila et al., 2017) for evidence of peptides produced during the MZT. We restricted our analysis to coding sequence uniquely present in the zygotic isoforms, of which 167 zygotic isoforms (86.5%), had distinguishing sequences. Despite the substantially lower sensitivity of mass spectrometry to detect isoform differences, we observed peptides for 50 of the distinguishable zygotic isoforms (Figure 5C). Similarly, when we examined published Ribo-seq datasets (Eichhorn et al., 2016), we found evidence of translation for nearly two-thirds (66.5%) of the zygotic-specific isoforms (Figure 5D). Our inability to detect the remaining isoforms is likely partially explained by library coverage, lack of sufficient distinguishing CDS sequence, or translational repression of the zygotic isoform.

For instance, *PNUTS*, a protein phosphatase 1 (PP1) binding protein) regulates transcriptional elongation and transcription termination through modulation of PP1 phosphatase activity (Cortazar et al., 2019). The PP1 interaction motif (residues 720-726) is absent from the dominant zygotic isoform, and this isoform is translated in the time points following zygotic transcription (Figure 5E). The lack of this domain may alter RNA polymerase II phosphorylation and transcription dynamics (Ciurciu et al., 2013). Similarly, *CycE* also shows differences in maternal and zygotic isoforms. The zygotic transcript incorporates a distinct in-frame 5’UTR exon that alters the first 12 amino acid and truncates 107 amino acids from the N-terminus of the maternal isoform. Although the zygotic transcript had previously been characterized by *in situ* hybridization (Richardson et al., 1993), our analysis shows that the shortened zygotic transcript is translated. Given the tight control of the cell cycle during the MZT, the central role cyclin E plays in cell cycle progression, and the fact that *Cdk2*, a binding partner, also has different zygotic isoforms, this intriguing observation warrants future study.

## DISCUSSION

Annotations of maternal or zygotic genes in *D. melanogaster* have used a number of clever experimental tools, including deletion chromosomes, metabolic labeling, and SNP-mapping (De Renzis et al., 2007; Kwasnieski et al., 2019; Lott et al., 2011). These methods are powerful, but they can be harder to apply to other organisms with fewer genetic tools, even including other *Drosophilids*. Our motivation for this study was to develop an easy and robust computational method that could determine transcriptome dynamics during the MZT using only RNA-seq and so could be applied to a wide variety of organisms. Like others (Hwang et al., 2018; Lee et al., 2013), we reasoned that, based on the instability of introns, reads mapping to these regions could act as a proxy for transcription. Our computational innovation of assembling the maternal transcriptome based on pre-ZGA datasets and implementing a two-pass alignment approach to quantify intron signal was an improvement over previous methods. We also accounted for ambiguous read alignments, further improving estimates of intronic signals. We showed that this method was applicable to a number of animal MZT systems, including those like chicken that lack the types of genetic tools found in *Drosophila*. Moreover, conceptually, our method need not be restricted to the MZT because it can easily be applied to other dynamic processes that may be less amenable to experimental manipulation like metabolic labeling.

Nonetheless, there are caveats for our method. The first major requirement is that the genome of the animal of interest must be relatively well annotated. The second is that, unsurprisingly, this method requires introns within the gene to be present and annotated. For many model organisms, this requirement may not impose a major limitation, but our results with *D. melanogaster* highlight this caveat: our method failed to annotate experimentally identified ZGA genes that lack introns and that also make up a major population of genes initially transcribed in the embryo (De Renzis et al., 2007). The final major caveat arises from the fact that standard RNA-seq library preparations use oligo(dT) enrichment. Because the MZT is often associated with changes in poly(A)-tail length (Barkoff et al., 1998; Barkoff et al., 2000; Eichhorn et al., 2016; Gebauer et al., 1994), poly(A)-tail lengthening can lead to the spurious identification of maternally deposited mRNAs as zygotically transcribed. Because this caveat extends to all RNA quantification during the MZT based upon oligo(dT) enrichment, our recommendation is to use rRNA depletion whenever possible for these studies. An added benefit of rRNA depletion is that reads mapping to introns (which lack poly(A) tails) are also more abundant in these libraries, leading to easier classification of transcription.

Although our initial goal was to characterize zygotic transcription, our transcript-based method opened the door to investigating how transcript isoforms change during the MZT. In contrast to gene-level annotations, there has been relatively little work done on the topic of isoform changes, even in well-characterized model organisms like *Drosophila*. Previous studies have highlighted the role of alternative polyadenylation during ZGA, and a contemporaneous study has pointed to sex-specific alternative splicing during zygotic transcription (Ray et al., 2021). Our results extend these examples by fully characterizing the types of differences that exist between maternal and zygotic transcript isoforms. Indeed, zygotic isoforms can differ in promoter, poly(A) signal, and splicing—and usually a combination of all three. These complex differences between maternal and zygotic transcript isoforms mean that the two sets of transcripts are likely subject to different post-transcriptional regulation and, more often than not, encode a different protein isoform.

Our results raise a series of related questions. First, how do changes in the pre-mRNA processing machinery lead to these differences? Given that not all newly made transcripts in the embryo use different poly(A) signals or promoters or are spliced differently, what common regulatory mechanisms can explain those genes that do show changes? It is intriguing to note that recent work on splicing and poly(A) signal recognition can be affected by RNA polymerase speed (Cortazar et al., 2019), and that PNUTS, a key regulator of RNA pol II (Ciurciu et al., 2013), is itself controlled by alternative splicing. Exploring these issues is an important next step.

Second, what are the consequences of maternal and zygotic isoform differences? Changes in 5′ and 3′UTRs clearly have the potential to alter post-transcriptional regulation, and one possibility is that these differences allow selective regulation (such as mRNA clearance or translation) of the maternal, rather than zygotic, isoform. Our observation that zygotic isoforms often also differ in their coding region is an intriguing new level of complexity and provides a rationale as to how different translational control of the maternal and zygotic isoforms could have a biological impact. In this respect, *toc*, whose zygotic isoform has altered 5′UTR (which is predicted to be more amenable to translation) and shortened coding region, may be an interesting case study, especially given its known role in mitotic spindle formation and cellularization (Debec et al., 2001).

More generally, an important consideration is the extent to which the maternal protein isoform is expressed after ZGA. In other words, is the maternal protein isoform replaced by the zygotic one? How does this replacement affect gene function? These questions tap into the relatively unexplored area of protein decay during the MZT, and we suspect that biologically important shifts in gene function will be characterized by unstable maternal protein isoforms that hand off to well-expressed zygotic ones, enabling a change in protein activity. In this respect, our observations of changes in *CycE* are especially intriguing, given that its corresponding E3 ubiquitin ligase of Archipelago is also expressed during the MZT based on RNA-seq data (Graveley et al., 2011; Moberg et al., 2001). Finally, as we learn more about transcript diversity in the MZT, it will be important to take an evolutionary eye to these results and explore the common, and distinct, regulatory levers employed across the animal kingdom.

## ACKNOWLEDGEMENTS

We thank members of the Rissland lab for helpful and thoughtful discussions, and the RNA Bioscience Initiative Informatics Fellows for technical discussions. We are grateful to Dr. Hesselberth, Dr. Mukherjee, and Dr. Ramachandran for their feedback on this manuscript. This work was supported by NIH grants R35GM128680 (OSR) and the RNA Bioscience Initiative, University of Colorado School of Medicine. KR was supported as an Informatics Fellow of the RNA Bioscience Initiative, University of Colorado School of Medicine.

## MATERIALS AND METHODS

### Drosophila fly stocks

Fly stocks (*w1118*) were maintained in a 25°C incubator with 65% humidity.

### RNA extraction from embryos

Embryos were collected at various time points post-egg laying, dechorionated with bleach, and washed with 0.1% Triton X-100. Embryos were then homogenized in lysis Buffer B (150 mM KCl, 20 mM HEPES-KOH pH 7.4, 1 mM MgCl_2_, 1mM DTT, complete mini EDTA-free protease inhibitors), and were clarified at 15,000 rpm, 4°C for 15 minutes. The supernatant was stored at −80°C. RNA was extracted using TRI-reagent, according to the manufacturer’s instructions.

### Nanopore sequencing

Nanopore libraries were prepared according to the Direct RNA sequencing protocol (SQK-RNA002). Because the lengths of poly(A) tails change during the MZT, total RNA was used in place of oligo(dT)-selected RNA. Libraries were sequenced on a FLO-MIN106 flow cell and minION sequencing device. Base calling was performed using Guppy (v. 3.1.5+781ed57) using default settings. Sequencing reads from n = 2 replicate libraries from each time point were aligned to the Drosophila transcriptome and independently to the genome using minimap2 (Li, 2018). Transcript abundance was quantified using Salmon and read counts from each time point were combined. Genome alignments were used for visualizing read coverage with each read assigned to a transcript based on the transcriptome alignment. Raw sequencing data is available from GEO (GSE183975).

### Intron quantification pipeline

Sequencing reads were preprocessed to remove adapter sequences and trim low quality bases using cutadapt (-q 10, v.1.16, (Martin, 2011)). EisaR (Soneson et al., 2021) was used to generate a transcriptome FASTA file containing spliced transcripts and intronic sequences with 40 bp of flanking exonic sequence. Samples from the earliest timepoint, prior to zygotic transcription (*e.g*., 0-1 hr A.E.L), were aligned to the intron supplemented transcriptome using bowtie2 (v.2.3.2, (Langmead and Salzberg, 2012)) with parameters set to report up to 250 alignments (-k 250). Read coverage tracks were generated using bedtools (2.26.0). Intronic sequences with read alignments were next masked to an ambiguous nucleotide (N) to avoid alignment and subsequent quantification of intronic signals derived from RNAs produced prior to zygotic transcription. Introns with > 80% masked sequence or less than 25 unmasked nucleotides were discarded. All samples were then aligned to the masked transcriptome followed by transcript and intron abundance quantification using salmon in alignment mode (v.1.1.0, (Patro et al., 2017)). Transcript level abundance estimates were then aggregated to generate gene-level estimates of spliced and intronic sequences using tximport (Soneson et al., 2015). Differential expression between maternal and zygotic stages was conducted using DESeq2 (Love et al., 2014) with an adjusted p-value < 0.05 considered significant.

Genes were next classified into four categories: strictly maternal, ambiguous maternal, purely zygotic, and maternal and zygotic. Genes with zygotic transcription were required to have significance differentiation expression (adjusted p-value < 0.05 and > 2-fold increase) in either exonic or intronic sequence. Maternal genes were required to be expressed at >1 TPM. Purely zygotic genes had no detectable maternal contribution (<1 TPM). Purely maternal genes were further subdivided into strictly maternal and ambiguous maternal. Strictly maternal genes had intronic signals <1 TPM throughout the entire time course or were mono-exonic, whereas ambiguous maternal genes had non-significant (> 0.05) exonic or intronic increases, with intronic TPM rising to >1 TPM in the time course.

Genes with zygotic transcription were assigned to the first time point of detectable expression based on the first time point with intronic levels >1 TPM. Purely zygotic genes with no intronic signal were assigned based on the first time point with >1 TPM of exonic signal.

The pipeline is available as a snakemake workflow provided at https://github.com/rnabioco/mzt-introns. Unless otherwise stated the analysis steps described were conducted in R (v. 4.0.3).

### Comparison to traditional intron quantification pipeline

To benchmark our method we applied the intron quantification strategies employed in previous studies to the rRNA-depleted *Drosophila* embryogenesis dataset (Hwang et al., 2018; Lee et al., 2013). Following adapter trimming, reads were aligned to the genome using STAR (Dobin et al., 2013). Gene-level exonic and intronic SAF gene annotation files were generated. Intronic regions with read alignments from the earliest maternal sample (0-1 AEL) were excluded. featureCounts was used to quantify uniquely aligned reads overlapping exons (--fracOverlap 1) or introns (--minOverlap 10) separately (Liao et al., 2014). Reads that aligned to both exonic and intronic features were ignored.

### Transcript isoform analysis

DEXSeq (v. 1.38.0) was used to identify differential transcript usage in pre (0-1 hr and 1-2 hr AEL) versus post major wave (2-3, 3-4, and 4-5 hr AEL) samples. Per transcript p-values were aggregated to identify genes with alterations in transcript usage with an adjusted p-value < 0.05 considered significant. For each significant gene, the highest abundance transcript in the pre or post major-wave samples was selected as the representative maternal or zygotic isoform respectively. The representative zygotic transcript was required to account for at least 10% of the expressed transcripts, have a transcript level adjusted p-value < 0.05, and increase in abundance in the post major wave samples.

### Analysis of published datasets

Previous classifications of zygotic *Drosophila* genes were taken from supplemental data files of published studies (De Renzis et al., 2007; Kwasnieski et al., 2019; Lott et al., 2011). Zelda binding was examined using supplemental data from a previously published Chip-Seq dataset (Harrison et al., 2011). Peptide level mass spectrometry data from *Drosophila* embryogenesis was obtained from Flybase (FBrf0241309 and 28381612 (Casas-Vila et al., 2017)). Protein sequence unique to zygotic isoforms was matched against peptides present in the mass spectrometry data. Poly(A)+ RNA-seq and ribosome profiling from *Drosophila* oogenesis and early embryogenesis were obtained from GEO dataset GSE83616 (Eichhorn et al., 2016). Ribosome profiling data was quantified by alignment to the genome using STAR, converted to BigWig read coverage files, and quantified by counting the number of alignments overlapping CDS regions unique to zygotic isoforms. rRNA-depleted RNA-seq data from *Drosophila melanogaster*, poly(A)+ RNA-seq data from *Gallus gallus*, poly(A)+ and rRNA-depleted RNA-seq data from *Xenopus tropicalis*, and poly(A)+ RNA-seq data from *Danio rerio*, were obtained from GEO datasets GSE98106, GSE86592, GSE65785, and dataset PRJEB12982 from the European Nucleotide Archive, respectively (Hwang et al., 2018; Owens et al., 2016; Wang et al., 2017; White et al., 2017). Genome and transcript references used in this study: *Drosophila melanogaster* (Ensembl BDGP6 release 84) *Gallus gallus* (Ensembl gallus-gallus-5 release 94), *Xenopus tropicalis* (UCSC xenTro9) and *Danio Rerio* (UCSC danRer10).

## SUPPLEMENTAL MATERIAL

**Supplemental Figure 1. Intronic read densities for maternal or zygotic genes.** Intron coverage of RNA-seq libraries generated using oligo(dT) selection or rRNA-depletion. Gene sets were derived from Lott et al., 2011.

**Supplemental Figure 2. Impact on maternal classifications by filtering for maximal intron TPM values and p-values from differential expression testing.** Maternal transcripts were subdivided into strictly maternal and ambiguous maternal, due to a subset of maternal transcripts exhibiting non-significant intronic increases during ZGA. The selected cutoffs for strictly maternal mRNAs were intronic levels < 1 TPM and an adjusted p-value from intronic differential expression testing > 0.05.

**Supplemental Figure 3. Examination of genes uniquely detected by the traditional alignment method. (** A) Summary of explanations for the lack of detection of zygotic genes by our pipeline. (B) Mean TPM values for genes missed by our pipeline. (C) Number of gene-level exonic or intronic features detectable by our pipeline or the traditional pipeline. (D) Mean TPM distributions for all genes throughout embryogenesis.

**Supplemental Figure 4. Previous intron-read mapping methods identify fewer zygotic transcripts.** A Venn diagram showing the overlap between zygotic transcripts identified by counting reads mapping to introns and those identified by different experimental methods.

**Supplemental Table 1. Zygotic and maternal expressed gene classifications.**

**Supplemental Table 2. Transcript-level classifications for maternal and zygotic expression.**

**Supplemental Table 3. GO term enrichments for genes with different maternal and zygotic isoforms.**

## REFERENCES

Barkoff, A., Ballantyne, S. and Wickens, M. (1998). Meiotic maturation in Xenopus requires polyadenylation of multiple mRNAs. EMBO J. 17, 3168–3175.

Barkoff, A. F., Dickson, K. S., Gray, N. K. and Wickens, M. (2000). Translational control of cyclin B1 mRNA during meiotic maturation: coordinated repression and cytoplasmic polyadenylation. Dev. Biol. 220, 97–109.

Benoit, B., He, C. H., Zhang, F., Votruba, S. M., Tadros, W., Westwood, J. T., Smibert, C. A., Lipshitz, H. D. and Theurkauf, W. E. (2009). An essential role for the RNA-binding protein Smaug during the Drosophila maternal-to-zygotic transition. Development 136, 923–932.

Bushati, N., Stark, A., Brennecke, J. and Cohen, S. M. (2008). Temporal reciprocity of miRNAs and their targets during the maternal-to-zygotic transition in Drosophila. Curr. Biol. 18, 501–506.

Casas-Vila, N., Bluhm, A., Sayols, S., Dinges, N., Dejung, M., Altenhein, T., Kappei, D., Altenhein, B., Roignant, J.-Y. and Butter, F. (2017). The developmental proteome of Drosophila melanogaster. Genome Res. 27, 1273–1285.

Chen, L., Dumelie, J. G., Li, X., Cheng, M. H., Yang, Z., Laver, J. D., Siddiqui, N. U., Westwood, J. T., Morris, Q., Lipshitz, H. D., et al. (2014). Global regulation of mRNA translation and stability in the early Drosophila embryo by the Smaug RNA-binding protein. Genome Biol. 15, R4.

Ciurciu, A., Duncalf, L., Jonchere, V., Lansdale, N., Vasieva, O., Glenday, P., Rudenko, A., Vissi, E., Cobbe, N., Alphey, L., et al. (2013). PNUTS/PP1 regulates RNAPII-mediated gene expression and is necessary for developmental growth. PLoS Genet. 9, e1003885.

Cortazar, M. A., Sheridan, R. M., Erickson, B., Fong, N., Glover-Cutter, K., Brannan, K. and Bentley, D. L. (2019). Control of RNA Pol II Speed by PNUTS-PP1 and Spt5 Dephosphorylation Facilitates Termination by a “Sitting Duck Torpedo” Mechanism. Mol. Cell 76, 896–908.e4.

De Renzis, S., Elemento, O., Tavazoie, S. and Wieschaus, E. F. (2007). Unmasking activation of the zygotic genome using chromosomal deletions in the Drosophila embryo. PLoS Biol. 5, e117.

Debec, A., Grammont, M., Berson, G., Dastugue, B., Sullivan, W. and Couderc, J. L. (2001). Toucan protein is essential for the assembly of syncytial mitotic spindles in Drosophila melanogaster. Genesis 31, 167–175.

Dobin, A., Davis, C. A., Schlesinger, F., Drenkow, J., Zaleski, C., Jha, S., Batut, P., Chaisson, M. and Gingeras, T. R. (2013). STAR: ultrafast universal RNA-seq aligner. Bioinformatics 29, 15–21.

Edgar, B. A. and Schubiger, G. (1986). Parameters controlling transcriptional activation during early Drosophila development. Cell 44, 871–877.

Eichhorn, S. W., Subtelny, A. O., Kronja, I., Kwasnieski, J. C., Orr-Weaver, T. L. and Bartel, D. P. (2016). mRNA poly(A)-tail changes specified by deadenylation broadly reshape translation in Drosophila oocytes and early embryos. Elife 5, 714.

Gaidatzis, D., Burger, L., Florescu, M. and Stadler, M. B. (2015). Analysis of intronic and exonic reads in RNA-seq data characterizes transcriptional and post-transcriptional regulation. Nat. Biotechnol. 33, 722–729.

Gebauer, F., Xu, W., Cooper, G. M. and Richter, J. D. (1994). Translational control by cytoplasmic polyadenylation of c-mos mRNA is necessary for oocyte maturation in the mouse. EMBO J. 13, 5712–5720.

Ghosh, S., Wang, Y., Cook, J. A., Chhiba, L. and Vaughn, J. C. (2013). A molecular, phylogenetic and functional study of the dADAR mRNA truncated isoform during Drosophila embryonic development reveals an editing-independent function. Open J Anim Sci 3, 20–30.

Giraldez, A. J., Cinalli, R. M., Glasner, M. E., Enright, A. J., Thomson, J. M., Baskerville, S., Hammond, S. M., Bartel, D. P. and Schier, A. F. (2005). MicroRNAs regulate brain morphogenesis in zebrafish. Science 308, 833–838.

Graveley, B. R., Brooks, A. N., Carlson, J. W., Duff, M. O., Landolin, J. M., Yang, L., Artieri, C. G., van Baren, M. J., Boley, N., Booth, B. W., et al. (2011). The developmental transcriptome of Drosophila melanogaster. Nature 471, 473–479.

Guo, X., Macleod, G. T., Wellington, A., Hu, F., Panchumarthi, S., Schoenfield, M., Marin, L., Charlton, M. P., Atwood, H. L. and Zinsmaier, K. E. (2005). The GTPase dMiro is required for axonal transport of mitochondria to Drosophila synapses. Neuron 47, 379–393.

Harrison, M. M., Botchan, M. R. and Cline, T. W. (2010). Grainyhead and Zelda compete for binding to the promoters of the earliest-expressed Drosophila genes. Dev. Biol. 345, 248–255.

Harrison, M. M., Li, X.-Y., Kaplan, T., Botchan, M. R. and Eisen, M. B. (2011). Zelda binding in the early Drosophila melanogaster embryo marks regions subsequently activated at the maternal-to-zygotic transition. PLoS Genet. 7, e1002266.

Heyn, P., Kircher, M., Dahl, A., Kelso, J., Tomancak, P., Kalinka, A. T. and Neugebauer, K.M. (2014). The earliest transcribed zygotic genes are short, newly evolved, and different across species. Cell Rep. 6, 285–292.

Hwang, Y. S., Seo, M., Kim, S. K., Bang, S., Kim, H. and Han, J. Y. (2018). Zygotic gene activation in the chicken occurs in two waves, the first involving only maternally derived genes. Elife 7,.

Kwasnieski, J. C., Orr-Weaver, T. L. and Bartel, D. P. (2019). Early genome activation in Drosophila is extensive with an initial tendency for aborted transcripts and retained introns. Genome Res. 29, 1188–1197.

La Manno, G., Soldatov, R., Zeisel, A., Braun, E., Hochgerner, H., Petukhov, V., Lidschreiber, K., Kastriti, M. E., Lönnerberg, P., Furlan, A., et al. (2018). RNA velocity of single cells. Nat. Biotechnol. 560, 494–498.

Langmead, B. and Salzberg, S. L. (2012). Fast gapped-read alignment with Bowtie 2. Nat. Methods 9, 357–359.

Laver, J. D., Li, X., Ray, D., Cook, K. B., Hahn, N. A., Nabeel-Shah, S., Kekis, M., Luo, H., Marsolais, A. J., Fung, K. Y., et al. (2015a). Brain tumor is a sequence-specific RNA-binding protein that directs maternal mRNA clearance during the Drosophila maternal-to-zygotic transition. Genome Biol. 16, 94.

Laver, J. D., Marsolais, A. J., Smibert, C. A. and Lipshitz, H. D. (2015b). Regulation and Function of Maternal Gene Products During the Maternal-to-Zygotic Transition in Drosophila. Curr. Top. Dev. Biol. 113, 43–84.

Lécuyer, E., Yoshida, H., Parthasarathy, N., Alm, C., Babak, T., Cerovina, T., Hughes, T. R., Tomancak, P. and Krause, H. M. (2007). Global analysis of mRNA localization reveals a prominent role in organizing cellular architecture and function. Cell 131, 174–187.

Lee, M. T., Bonneau, A. R., Takacs, C. M., Bazzini, A. A., DiVito, K. R., Fleming, E. S. and Giraldez, A. J. (2013). Nanog, Pou5f1 and SoxB1 activate zygotic gene expression during the maternal-to-zygotic transition. Nature 503, 360–364.

Li, H. (2018). Minimap2: pairwise alignment for nucleotide sequences. Bioinformatics 34, 3094–3100.

Liang, H.-L., Nien, C.-Y., Liu, H.-Y., Metzstein, M. M., Kirov, N. and Rushlow, C. (2008). The zinc-finger protein Zelda is a key activator of the early zygotic genome in Drosophila. Nature 456, 400–403.

Liao, Y., Smyth, G. K. and Shi, W. (2014). featureCounts: an efficient general purpose program for assigning sequence reads to genomic features. Bioinformatics 30, 923–930.

Lott, S. E., Villalta, J. E., Schroth, G. P., Luo, S., Tonkin, L. A. and Eisen, M. B. (2011). Noncanonical compensation of zygotic X transcription in early Drosophila melanogaster development revealed through single-embryo RNA-seq. PLoS Biol. 9, e1000590.

Love, M. I., Huber, W. and Anders, S. (2014). Moderated estimation of fold change and dispersion for RNA-seq data with DESeq2. Genome Biol. 15, 550.

Lugowski, A., Nicholson, B. and Rissland, O. S. (2018). DRUID: a pipeline for transcriptome-wide measurements of mRNA stability. RNA 24, 623–632.

Martin, M. (2011). Cutadapt removes adapter sequences from high-throughput sequencing reads. EMBnet.journal 17, 10–12.

McDaniel, S. L., Gibson, T. J., Schulz, K. N., Fernandez Garcia, M., Nevil, M., Jain, S. U., Lewis, P. W., Zaret, K. S. and Harrison, M. M. (2019). Continued Activity of the Pioneer Factor Zelda Is Required to Drive Zygotic Genome Activation. Mol. Cell 74, 185–195.e4.

Moberg, K. H., Bell, D. W., Wahrer, D. C., Haber, D. A. and Hariharan, I. K. (2001). Archipelago regulates Cyclin E levels in Drosophila and is mutated in human cancer cell lines. Nature 413, 311–316.

Nien, C.-Y., Liang, H.-L., Butcher, S., Sun, Y., Fu, S., Gocha, T., Kirov, N., Manak, J. R. and Rushlow, C. (2011). Temporal coordination of gene networks by Zelda in the early Drosophila embryo. PLoS Genet. 7, e1002339.

Owens, N. D. L., Blitz, I. L., Lane, M. A., Patrushev, I., Overton, J. D., Gilchrist, M. J., Cho, K. W. Y. and Khokha, M. K. (2016). Measuring Absolute RNA Copy Numbers at High Temporal Resolution Reveals Transcriptome Kinetics in Development. Cell Rep. 14, 632–647.

Patro, R., Duggal, G., Love, M. I., Irizarry, R. A. and Kingsford, C. (2017). Salmon provides fast and bias-aware quantification of transcript expression. Nat. Methods 14, 417–419.

Ray, M., Conard, A. M., Urban, J. and Larschan, E. (2021). Sex-specific transcript diversity is regulated by a maternal pioneer factor in early Drosophila embryos. bioRxiv 2021.03.18.436074.

Richardson, H. E., O’Keefe, L. V., Reed, S. I. and Saint, R. (1993). A Drosophila G1-specific cyclin E homolog exhibits different modes of expression during embryogenesis. Development 119, 673–690.

Rothe, M., Pehl, M., Taubert, H. and Jäckle, H. (1992). Loss of gene function through rapid mitotic cycles in the Drosophila embryo. Nature 359, 156–159.

Shermoen, A. W. and O’Farrell, P. H. (1991). Progression of the cell cycle through mitosis leads to abortion of nascent transcripts. Cell 67, 303–310.

Soneson, C., Love, M. I. and Robinson, M. D. (2015). Differential analyses for RNA-seq: transcript-level estimates improve gene-level inferences. F1000Res. 4, 1521.

Soneson, C., Srivastava, A., Patro, R. and Stadler, M. B. (2021). Preprocessing choices affect RNA velocity results for droplet scRNA-seq data. PLoS Comput. Biol. 17, e1008585.

Tadros, W., Goldman, A. L., Babak, T., Menzies, F., Vardy, L., Orr-Weaver, T., Hughes, T. R., Westwood, J. T., Smibert, C. A. and Lipshitz, H. D. (2007). SMAUG is a major regulator of maternal mRNA destabilization in Drosophila and its translation is activated by the PAN GU kinase. Dev. Cell 12, 143–155.

Ulitsky, I., Shkumatava, A., Jan, C. H., Subtelny, A. O., Koppstein, D., Bell, G. W., Sive, H. and Bartel, D. P. (2012). Extensive alternative polyadenylation during zebrafish development. Genome Res. 22, 2054–2066.

Vastenhouw, N. L., Cao, W. X. and Lipshitz, H. D. (2019). The maternal-to-zygotic transition revisited. Development 146, dev161471–20.

Wang, M., Ly, M., Lugowski, A., Laver, J. D., Lipshitz, H. D., Smibert, C. A. and Rissland, O. S. (2017). ME31B globally represses maternal mRNAs by two distinct mechanisms during the Drosophila maternal-to-zygotic transition. Elife 6, 233.

White, R. J., Collins, J. E., Sealy, I. M., Wali, N., Dooley, C. M., Digby, Z., Stemple, D. L., Murphy, D. N., Billis, K., Hourlier, T., et al. (2017). A high-resolution mRNA expression time course of embryonic development in zebrafish. Elife 6,.

